# Highly Efficient Coreactant-Free Electrochemiluminescence Sensing Platform Using Novel Microfabricated Multiplexed Entwined Spiral Microelectrodes for Point-of-Care Applications

**DOI:** 10.1101/2025.11.03.686208

**Authors:** Somayyeh Bozorgzadeh, Richard Murray, Hassan Hamidi, Ian Seymour, Daniela Iacopino, Alan O’Riordan

## Abstract

Luminol-based Electrochemiluminescence (ECL) generates weak signals in neutral media and typically requires H_2_O_2_ as a coreactant. However, H_2_O_2_ instability and the need for on-site addition hinder real-time diagnostic applications. Compact integrated sensing platforms are ideal for point-of-care (POC) testing due to portability, low sample requirements, and multiplexing. However, as sensor dimensions decrease, the low light emission issue in ECL becomes severe.

We introduce a novel fully integrated miniaturized silicon device consisting of three sensors comprised of unique entwined micro-spiral electrodes in a generator-collector configuration. This enables highly sensitive, coreactant-free multiplexed sensing at physiological pH by accelerating in-situ reactive oxygen species production and boosting ECL intensity. Systematic optimization of electrode geometry (gaps and widths) yields an 11-fold improvement in ECL signal compared to a single-electrode setup, as well as excellent reproducibility and stability. In addition to Trolox and H_2_O_2_ detection, the platform demonstrates multiplex immunosensing through selective functionalization of the collector electrodes with chitosan nanocomposites, followed by Protein A/G-assisted immobilization of anti-IgG antibodies with peptide-based antifouling. The immunosensors exhibit high analytical performance (linear range: 0.001-100 pg·mL^−1^, detection limit: 0.8 fg·mL^−1^) with excellent reproducibility and reliability in 25% fetal bovine serum, highlighting the platform’s potential for sensitive, coreactant-free multi-analyte POC diagnostics.

## 1. Introduction

Electrochemiluminescence (ECL) has emerged as a powerful analytical technique, combining the advantages of both electrochemistry and chemiluminescence spectroscopy. It offers accurate spatial and temporal potential control, a simple operational setup, a wide dynamic concentration range, high sensitivity, and a low background signal.^[1,2]^ Therefore, ECL has attracted significant attention for its merits in a wide range of applications, including bioanalysis, food, environmental monitoring, immunoassays and clinical testing.^[3–8]^

However, despite the growing demand for highly sensitive point-of-care/use (POC/U) testing devices, particularly for on-site early diagnostics, the practical performance of ECL systems is still limited due to insufficient signal intensity, the requirement for high concentrations of coreactant reagents, the limited photostability of the luminophores, and interference from complex biological samples, which restrict their sensitivity and reliability in real-world applications.^[9–11]^ To address these challenges, various strategies have been explored to improve ECL performance, including the incorporation of novel nanomaterials to boost ECL signal intensity,^[12–14]^ the synthesis of highly efficient luminophores,^[15–17]^ the use of coreactant accelerators,^[18–22]^ and the development of highly effective antibiofouling interfaces in the structure of biosensors.^[23–28]^

Besides portability, ease of detection, multiplexed detection capabilities, cost-effectiveness, and reagentless ECL system within a compact, miniaturized platform are the critical factors for the development of field-deployable ECL-based POC systems.^[9,10,29]^ Although smartphone integration offers significant potential for improving portability and reducing cost in on-site monitoring, the sensitivity of phone-based ECL systems remains limited by the low sensitivity of smartphone camera.^[30–36]^ To overcome this issue, large electrodes and the high amounts of coreactant are often employed, which in turn restrict miniaturization, multiplexing, and portability. To date, no ECL system has successfully met all the essential criteria for POC testing. Consequently, ongoing research is focused on overcoming these individual limitations to develop fully functional, sensitive, and compact ECL based POC testing devices suitable for practical applications.^[9–11,37]^

Among various ECL luminophores, luminol is one of the most efficient reagents due to its low oxidation potential, low toxicity, high ECL efficiency, and cost-effectiveness. However, its ECL signal is weak under physiological conditions (pH 7.4), requiring enhancement through the external addition of hydrogen peroxide (H_2_O_2_) coreactant.^[38]^ However, H_2_O_2_ is unstable and must be added at the time of analysis, which makes it unsuitable for on-site and real-time diagnostics. Therefore, one promising step toward achieving a reagent less ECL system is the development of coreactant-free ECL systems by in-situ electrochemical generation of reactive oxygen species (ROSs) *via* dissolved oxygen reduction reaction (ORR). These ROSs can react with luminol radical anions, the product of luminol electro-oxidation, leading to strong and constant ECL emission.^[22,39,40]^ In-situ ROSs generation can be achieved either by applying two step potentials at a single working electrode (WE), for instance first by applying a reduction potential followed by a positive oxidation potential to oxidize luminol, or by simultaneously applying both potentials to dual WEs in generator-collector (GC) configuration.^[41,42]^ The first approach is more common and has been used with large electrodes. Consequently, most research has focused on developing novel nanostructured materials that accelerate ROSs production on a single WE rather than designing dual WEs, which can generate ROSs synchronously during luminol oxidation.^[40,43–46]^ Consequently, only two papers in the literature employing dual electrodes designs have been reported, both demonstrating a stronger and more stable ECL signal for luminol. This dual-WEs strategy has been applied to relatively large macroelectrodes: WE1 and WE2 (1 × 1 cm) in one design and WE1 (ITO glass, 1 × 5 cm) with WE2 (platinum net, 2 × 5 mm) in another. In both cases, while reducing the distance between the two WEs could enhance the ECL signal, the authors pointed out that they were unable to decrease it below 100 µm due to the risk of electrode short circuiting and asymmetrical current density distribution,^[41,42,47]^ highlighting the challenge of fabricating dual WEs for sensitive coreactant-free ECL applications. These challenges will become more difficult especially when applied to multiplexed platform.

To this end, recent advancements in the silicon microfabrication of robust and highly reproducible ultramicroelectrodes have opened the door to an exciting new area of fully integrated miniaturized multiplexed sensing platforms, which have been extensively used in the development of portable electrochemical based POC testing devices. Compared to traditional macroelectrodes, microscale electrodes offer significant advantages, including enhanced analyte mass transport *via* radial diffusion, reduced background noise due to their smaller surface area, and high current densities. So far, several microarrays with various shapes and designs have been developed for electrochemical analysis^[48–56]^

To bring silicon microfabricated multiplexed sensing platforms into the realm of coreactant-free ECL sensing systems, special design considerations and enhancements are required to overcome the low sensitivity of the ECL signal at microelectrodes compared to macroelectrodes. To address these challenges, we explored the design and fabrication of a fully integrated miniaturized silicon device consisting of three sensors, each comprising dual microelectrodes in an entwined spiral design, allowing multiplex detection. To the best of our knowledge, there are no reports on the design and application of a multi micro-sensing platform for coreactant free ECL applications.

Here, we examined seven unique entwined spiral microelectrodes by systematically varying electrode gap (G) and width (W) dimensions, enabling the tuning and exploration of the coreactant-free ECL response. This approach was compared with a single-electrode approach. Our results demonstrated that the spiral GC configuration significantly enhanced the ECL signal through facilitating simultaneous oxygen reduction at the generator electrode (GE) and luminol oxidation at the collector electrode (CE), offering a more effective solution for coreactant-free ECL sensing on a compact platform (**Scheme 1**). Following a comprehensive investigation of the influencing parameters and the underlying ECL mechanism, we evaluated the performance of this platform for Trolox and hydrogen peroxide detection. In addition, we investigated the electrode modifications for selective immobilization of antibodies on the CEs for immunoassay applications. This novel ECL platform exhibited remarkable sensitivity and potential for coreactant-free label free ECL immunoassay in a complex matrix, highlighting its suitability for multi-analyte detection within a single, integrated miniaturized sensing system. l

**Scheme 1:**
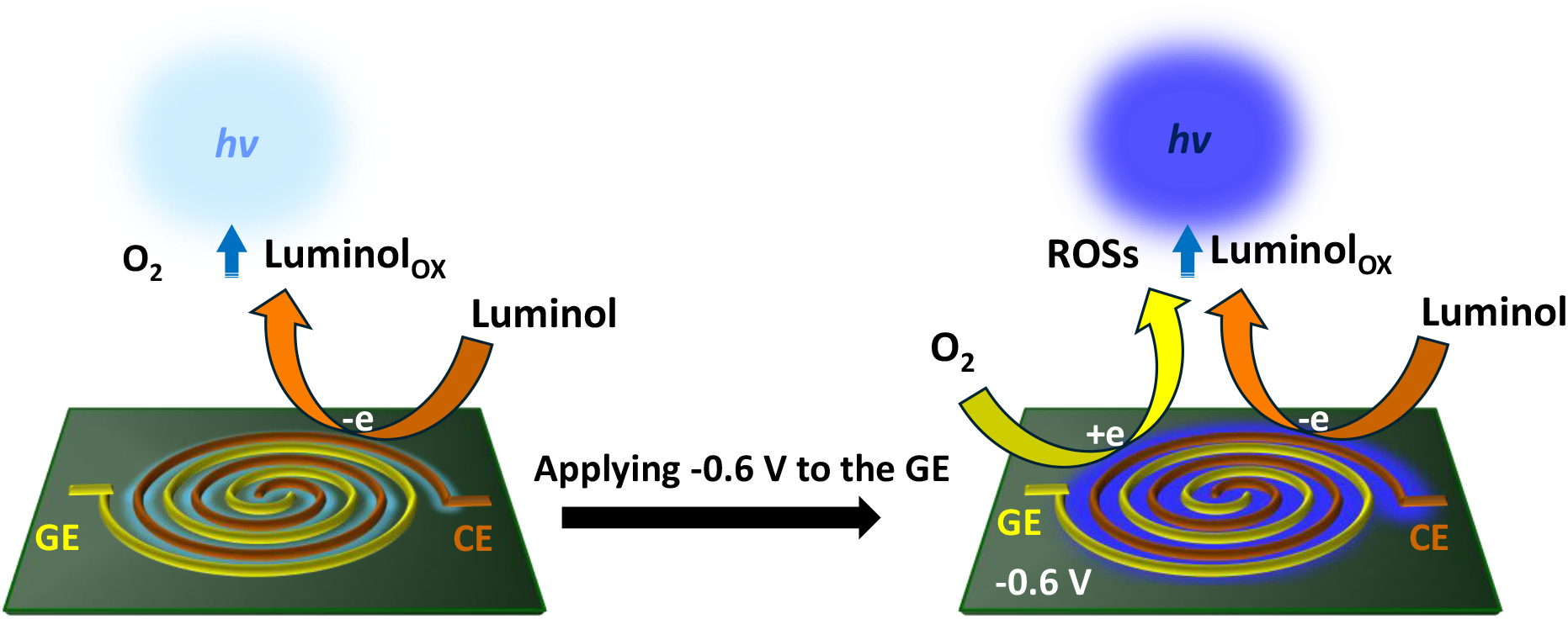
ECL mechanism pathway at entwined spiral microelectrodes without and with applying a negative bias to the GE. Significant ECL enhancement is observed when the -0.6 V vs on-chip platinum reference electrode is applied to the GE.

## 2. Experimental Section

### 2.1. Reagents

All chemicals were of analytical reagent grade and used without further purification. Potassium chloride (KCl), potassium ferrocyanide (K_4_[Fe(CN)_6_]), potassium ferricyanide (K_3_[Fe(CN)_6_]), acetone, phosphate buffer saline tablets (PBS, 0.01 M, pH7.4), gold(III)chloride trihydrate, ferrcene carboxylic acid, luminol, multi walled carbon nanotubes (MWCNTs), BSA, N-Hydroxysuccinimide (NHS) and N-(3-Dimethylaminopropyl)-N′-ethylcarbodiimide hydrochloride (EDC) were obtained from Sigma Aldrich (Dublin). Rabbit IgG protein, anti-Rabbit IgG and superblock were purchased from Thermofisher. Fetal Bovine Serum (FBS) was obtained from Generon The custom branched peptide (KE)4KE)2KEPPPPC was purchased from Merck (Dublin). Luminol stock solution (20 mM) was prepared by dissolving 0.0183 g of luminol in 5 mL NaOH 0.1M. All luminol solutions was diluted by 0.01 M PBS pH 7.4. A 5 mM Fe(CN)_6_]^3−/4−^ redox probe solution and 1mM ferrocenecarboxylic acid in 0.01 M PBS pH 7.4 and 0.1 M KCl was used for the electrochemical characterizations and redox cycling studies. All solutions were prepared with Milli-Q ultrapure water (18.2 MΩ cm ^−1^).

For the electrochemical characterizations and ECL studies on the bare gold microelectrodes, the device was cleaned firstly by washing in acetone and deionised water, 5 times each for 30 seconds, respectively. Once cleaned, the chip (device) was dried in a stream of nitrogen, placed in a 3D printed chip holder and connected to the potentiostat using a micro-SD connector and was further cleaned potentio-dynamically in 0.25 M KOH solution by running CVs from -1.2 -0.2 V versus on-chip Pt reference electrode.

### 2.2. Biosensor preparation

Chitosan (CS) stock solution (1%) was prepared by dissolving 1 g of chitosan in 100 ml deionised (DI) water containing 1% acetic acid. The solution was stirred overnight. Then, 500 µL of the chitosan (1%) solution was mixed with 500 µL MWCNTs dispersed solution in DIW (1 mg mL^-1^) and sonicated to ensure a well dispersed mixture. We used 50 mM HAuCl_4_ dissolved in 0.1 M KCl solution for gold nanostructured particles electrodeposition.

To develop ECL immunosensors on CEs, we chose chitosan, a biopolymer rich in amino functional groups, which provided the necessary binding sites for subsequent antibody immobilization and enabled selective electrodeposition on the CEs. First to increase the surface area, all six acetone-cleaned bare microelectrodes were connected and modified with gold nanostructures by applying a current of -1 mA for 5.0 s in 500 µL of 50 mM HAuCl_4_ dissolved in 0.1 M KCl solution. Subsequently, a chitosan-gold-MWCNTs nanocomposite hydrogel (CS-MWCNTs-Gold) was electrochemically deposited on top of the three CEs surfaces. The chitosan solution consisted of 500 µL of 0.1% (v/v) chitosan-MWCNTs and 2.5 mM HAuCl_4_. A voltage of -1.2 V (Vs Ag|AgCl|KCl _3M_) was applied to all three electrodes on a chip for 5 seconds. After deposition, for site directed immobilization, 50 µL protein A/G (50 µg/mL) mixed with 50 µL NHS:EDC (2:1). After activation of the carboxylic groups of protein A/G for 30 minutes, 2 µL of the mixture was dropped onto each electrode and stored overnight in a fridge. Electrodes were then washed with buffer and incubated with a branched peptide blocking agent for 1 hour to reduce non-specific binding. Finally, 2 µL of Anti-IgG (20 µg/mL) was applied to each electrode and incubated overnight and subsequently washed with PBS and used for IgG detection. The biosensors were kept in a fridge when was not being used.

### 2.3. Apparatus

Cyclic voltammetry (CV) studies were performed using an Multi AutoLab M101 (Utrecht, The Netherlands). ECL signals was recorded using a H11890-210 Photon Counting Head PMT module while a fixed potential and CV were simultaneously applied to the GE and CE in bipotential setup respectively. Both electrochemical and ECL studies was carried out in a 500 µL costum 3D-printed cell holder. **Figure S1** shows images of (A) a silicon chip device, (B) the 3D printed cell holder containing the device and a micro-SD connector, and (C) the ECL setup with a small PMT positioned vertically in front of the three sensors. Both a three and four on-device electrodes setup was employed for ECL measurements and electrochemical characterizations. A conventional three-electrode electrochemical setup was used for electrochemical modifications, a platinum counter electrode, and an external Ag|AgCl|KCl _3M_ reference electrode. On-chip Pt reference electrode also applied for ECL measurements and characterizations. Scanning electron microscopy (SEM) was performed with a Carl Zeiss Supra 40 Scanning Electron microscope for the characterizations of various designs and different modified electrodes. All measurements were performed at room temperature.

### 2.4. Silicon device multiplexed platform microfabrication

The silicon devices were manufactured at the Tyndall National Institute silicon microfabrication facility *via* photolithography, deposition, lift-off, and etching steps as reported previously ^[48,51–53]^ and illustrated in **Figure 1**. The process began with the thermal growth of a thick 300 nm silicon dioxide (SiO_2_) passivation layer on the substrate. To form the spiral sensing electrode, the pattern was transferred using metal mask-1 in the first photolithography step, followed by the deposition of a Titanium-Gold (Ti: Au; 10:150 nm) layer and a lift-off process. Metal mask-2 was then employed in the second photolithography step to create pin outs, interconnection tracks, and the counter and reference electrodes, employing a combination of Titanium, Gold, and Platinum (10 nm Ti/100 nm Au/10 nm Ti/100 nm Pt) deposited using electron beam evaporation, followed by another lift-off. A 500 nm thick silicon nitride (Si_3_N_4_) passivation layer was then applied to the wafers. Using a passivation mask, windows were etched to expose the underlying sensing electrodes, counter and reference electrodes, and contact pinouts. In the final step, the wafers were coated with a protective layer of photoresist before undergoing mechanical dicing, ensuring no damage to the chips (devices). (**Figure 1**). Each wafer consists of 28 devices with 7 designs, varying in width (W) and gap (G) on the micrometre scale and each device is named accordingly: W2G2, W2G5, W2G10, W2G20, W5G10, W10G10, and W20G10.

**Figure 1.**
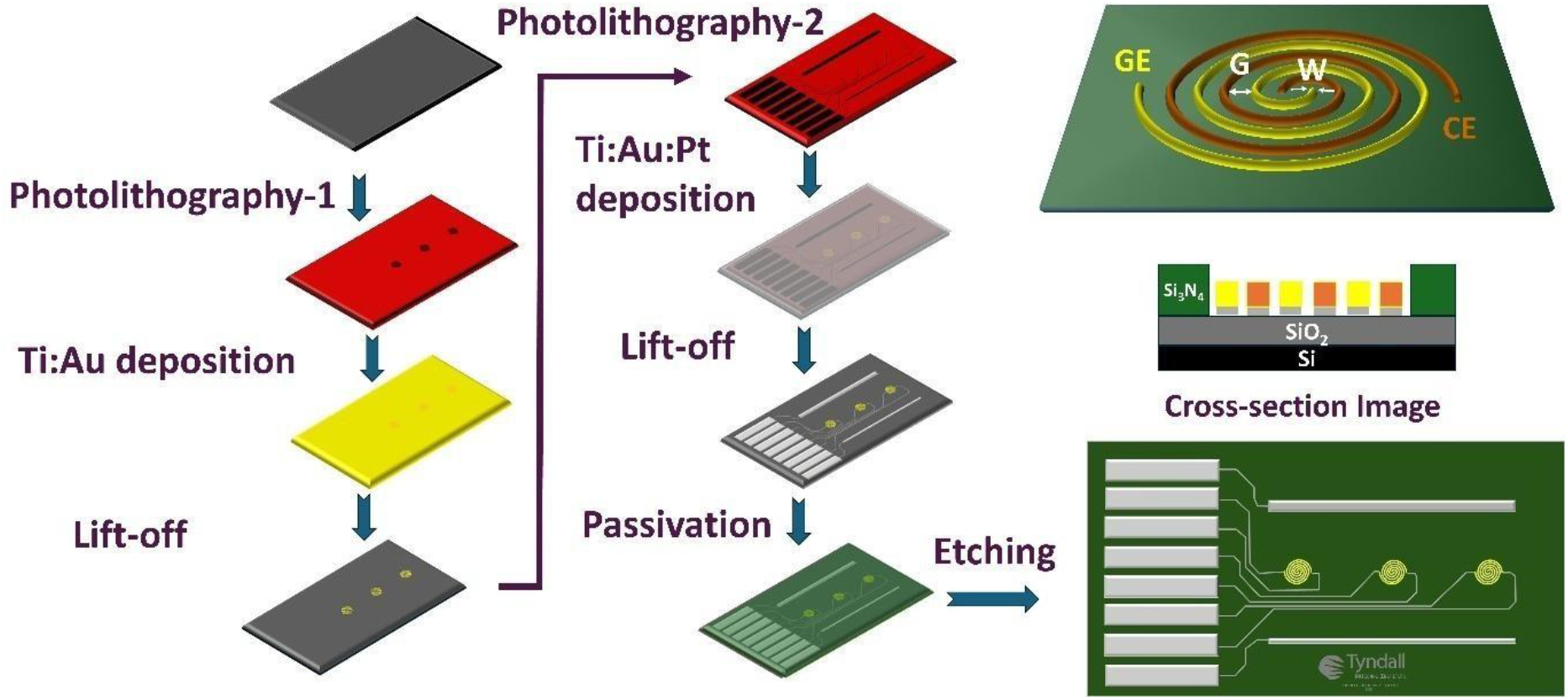
A Schematic illustration of gold entwined spiral microelectrodes fabrication on silicon device

## 3. Results and Discussion

### 3.1. SEM Characterizations

Figure 2 presents SEM images of seven designs of bare gold entwined spiral microelectrodes, demonstrating successful design and fabrication. As the gap between the spirals increased (**from A to D**) top row, the number of turns within each spiral decreased, which resulted in a reduced length and surface area. In comparison when the electrode width increased (**from D to G**) bottom row while keeping the gap size constant, the number of turns also decreased. However, overall, the larger electrode widths contribute to a higher overall surface area. Also, this study highlights the tunability of the electrode geometry and confirms the ability to fabricate entwined spiral microelectrodes with gaps and widths down to 2 µm using highly reproducible and precise technique.

### 3.2. Electrochemical Characterizations

Electrochemical characterizations of the seven designs were conducted using CVs of Fe(CN)_6_]^3−/4−^ redox couple on devices electrochemically cleaned with KOH solution. Figure 3A presents typical CVs at a W2G2 device at all three sensors GE1 to CE3 which are perfectly overlaid (RSD 0.7%), indicating that all six spiral electrodes are identical. Furthermore, we assessed the fabrication reproducibility by comparing anodic currents at three GEs and CEs across all three different W2G2 devices (**Figure 3B**, n=9). This analysis resulted in an RSD of 2.8 % for the GEs and 2.0% for the CEs, implying good consistency between inter and intra device fabrications. Also, further studies in all designs revealed that both cathodic and anodic peak currents decreased with increasing gap size from W2G2 to W2G10, due to a reduction in electrode surface area as the length of spiral shortens with wider gaps. However, at a fixed 10 µm gap, increasing the electrode width from W2G10 to W20G10 led to higher peak currents due to the greater surface area available for redox reactions (**Figure 3C** and **D**). These results emphasize the critical role of electrode geometry specifically gap and width on electrochemical performance.

**Figure 2.**
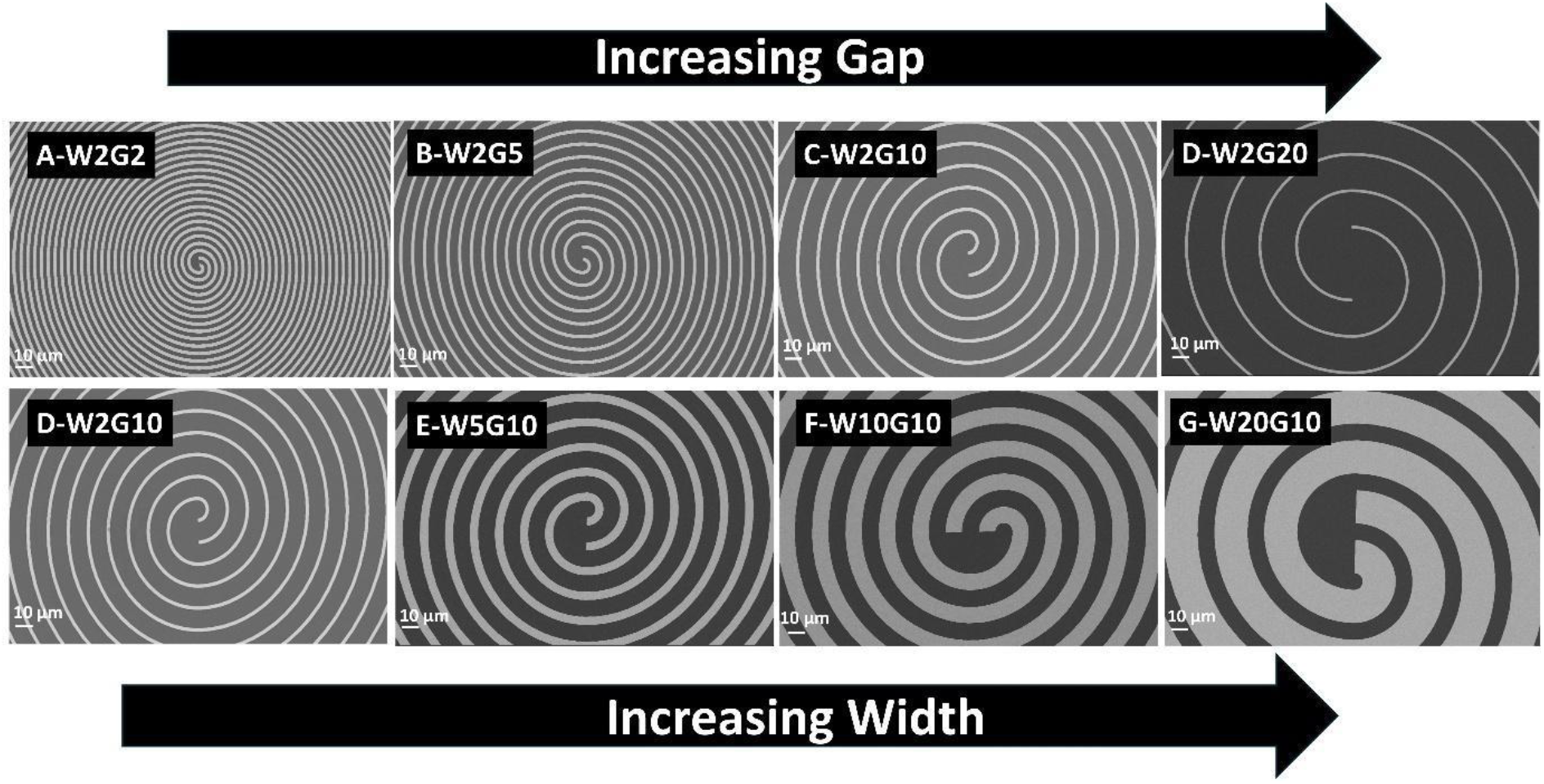
SEM images of seven designs of entwined spiral microelectrodes at 500x magnification.

**Figure 3.**
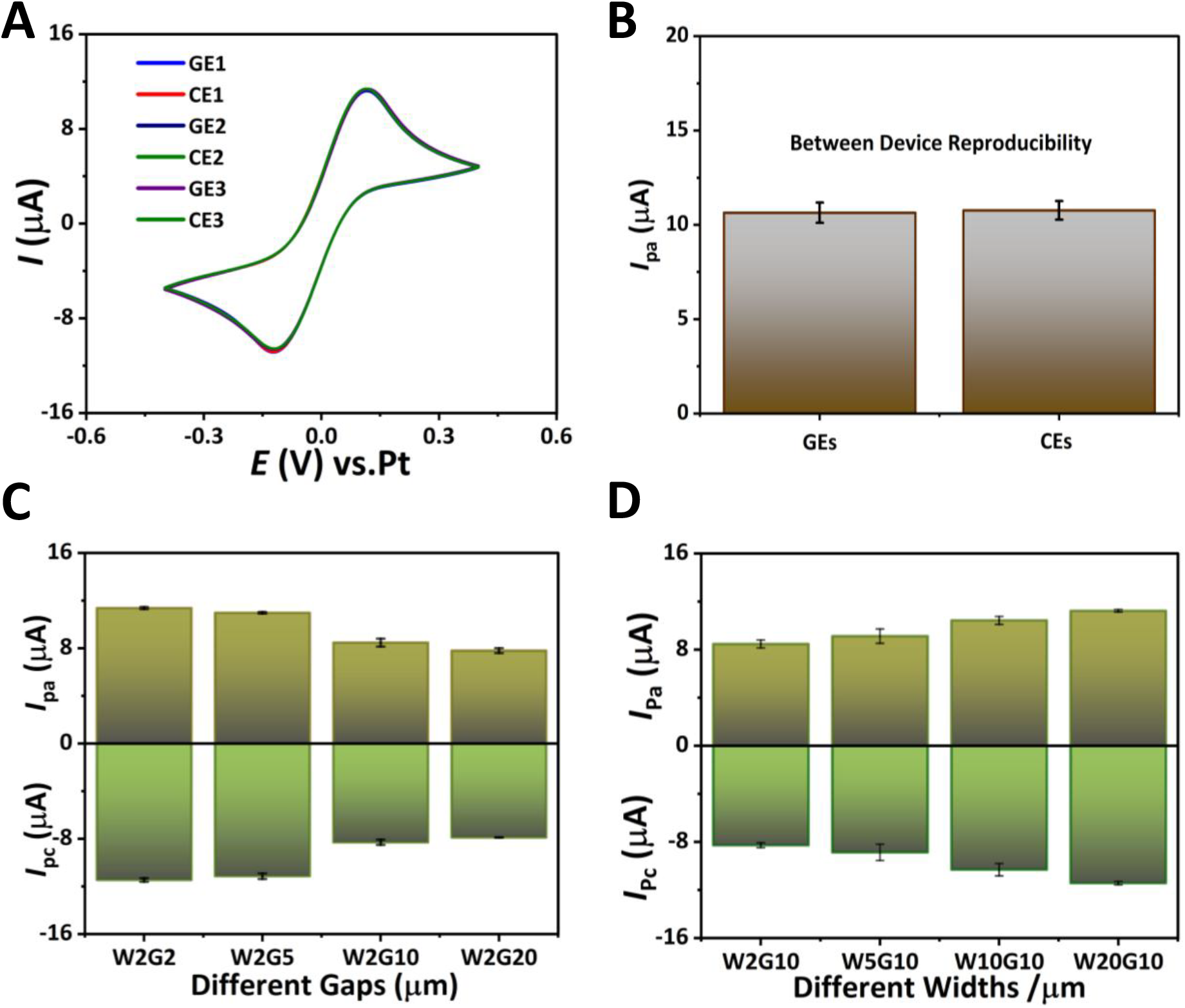
(A) CVs of 5 mM Fe(CN)_6_]^3−/4−^ redox probe at the W2G2 device (GE1 to CE3), (B) comparison of anodic peak currents among three independent W2G2 devices, the effect of varying (C) gaps and (D) widths on the anodic and cathodic peak currents at scan rate 0.1 V/s.

Next, redox cycling studies were performed to evaluate the collection efficiency of the GC designs throughout all seven designs (**Figure S2**). CVs of 1 mM ferrocene carboxylic acid in PBS (pH 7.4) were recorded at the GE, while a bias of 0.0 V versus on-chip Pt reference electrode was applied to the CE. The results revealed that increasing the gap between the GE and CE from 2 µm to 20 µm yielded a decrease in collection efficiency, from 93% for W2G2 to 56% for W2G20 (**Table S1**), due to the increased gap distance between GE and CE. Additionally, the effect of electrode width on redox cycling was examined using configurations W2G10, W5G10, W10G10, and W20G10. The results showed that collection efficiency remained almost unchanged across different widths (76 to 72 %), suggesting that gap size, rather than electrode width, is the dominant factor influencing diffusion and redox cycling efficiency.

### 3.3. Coreactant-free ECL behaviour of luminol

Firstly, ECL of luminol on the CE was investigated both with and without a negative potential bias being applied to a GE on W2G2 device. **Figure 4** compares the ECL response of 1 mM luminol in pH 7.4 under these conditions, while the potential of the CE simultaneously swept from 0 to 0.8 V versus on-chip Pt reference electrode. As illustrated in **Figure 4A**, only a very weak ECL signal was detected at the CE when not using the GE (single WE set-up). However, by applying -0.6 V to the GE (GC set-up strategy) significantly enhanced ECL signal of luminol by a factor of 11 (PBS pH 7.4).

**Figure 4.**
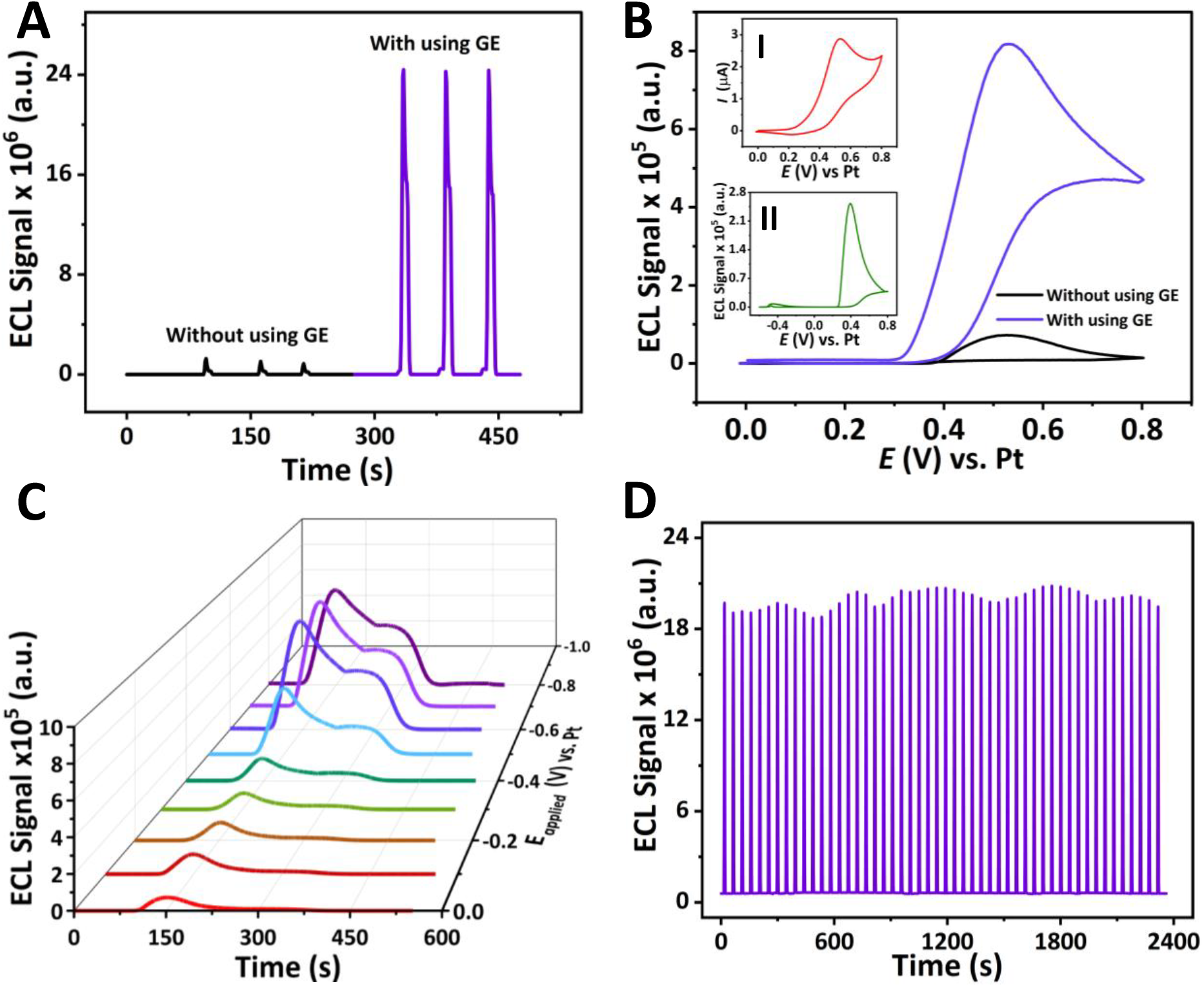
(A) ECL signal-time profile of luminol at the CE with and without using the GE; (B) ECL spectra (ECL signal versus potential). Inset I: CV of luminol in the range of 0-0.8 V and Inset II: ECL signal of luminol in the range of –0.6 to 0.8 V; (C) Effect of different applied potentials to the GE on the ECL signal; (D) Consecutive ECL signal recordings using GC setup. all in PBS (pH 7.4) containing 1 mM luminol.

**Figure 4B** presents the corresponding ECL responses (I_ECL_-E curves) and CV (inset I) of luminol at the CE in the potential range of 0.0 - 0.8 V using the W2G2 device, both with and without using the GE. To obtain ECL spectra, the ECL signals were recorded by PMT with a gate time of 33 milli seconds. The ECL signal of luminol reached its maximum at approximately 0.5 V, where luminol oxidation occurred, generating luminol radical anions for the subsequent chemiluminescent reaction. Notably, the ECL signal intensity of luminol increased 10.5-fold upon applying a negative potential to the GE, suggesting that negative potentials likely enhance the oxygen reduction reaction (ORR). This enhancement generates more reactive oxygen species (ROSs), which serve as co-reactants that react synergically with the produced luminol radical anions at the CE, significantly increasing the luminol ECL response.

Furthermore, we compared an alternative approach using single WE set-up where the potential was swept from -0.6 to 0.8 V at a CE (**Figure 4B**-**inset II**), instead of applying a constant negative potential at the GE. This potential sweep enhanced the signal approximately 2.5-fold compared to the condition without any applied negative potential to the GE (only applying 0.0 to 0.8 V at the CE), though the enhancement was significantly lower than that achieved with a constant -0.6 V potential at the GE. Additionally, we evaluated the effect of different applied potentials on a GE while sweeping the potential at the corresponding CE from 0.0 to 0.8 V. As shown in **Figure 4C**, the ECL signal increased with applying a negative potential from 0 to -0.6 V, reaching a plateau thereafter. Based on this observation, a potential of -0.6 V was therefore selected for further experiments across all different chip designs. Importantly, the stability and reproducibility of the luminol-ROSs system were analysed through 49 consecutive CV scans, yielding an RSD of 2.84% (**Figure 4D**). This result demonstrated the system’s stability and consistent performance over repeated measurements in a small volume (200 µL) of luminol solution.

In addition, as a multi-modal sensing platform, it is important to study the on-chip and between-chip reproducibility of ECL responses. To assess this, we selected three individual W2G2 devices and recorded the ECL signal with and without the use of the GE. For each study, three consecutive CVs were applied to the CE. The results in **Figure 5A** demonstrate the repeatability of the ECL signal across the three different devices. Without applying a negative potential to the GE, the ECL signal decayed over three consecutive CVs, resulting in low reproducibility (RSD: 20-25% between three WEs, with three measurements on each sensor).

**Figure 5.**
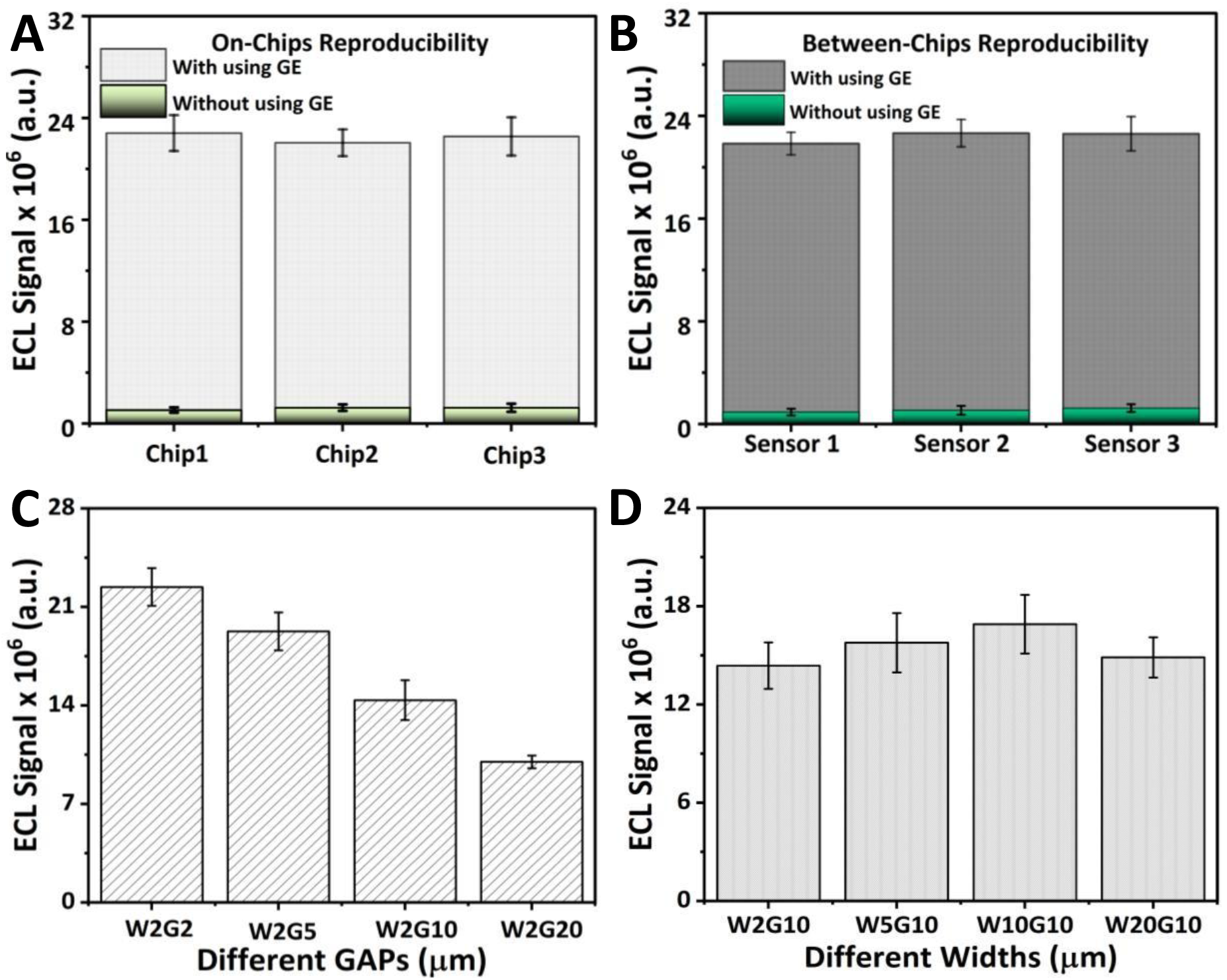
(A) comparison of on-chip; (B) between chip reproducibility of ECL signal for 1mM luminol at PBS pH 7.4 at W2G2 device; The effect of gaps (D) and widths (C) on ECL response of 1mM luminol while a potential -0.6 V was applied to the GE.

However, when a negative potential was applied to the GE, enhancing the ECL signal, the RSD of three continuous measurements decreased to 2-5%, showing a significant improvement in the repeatability of the multi-ECL platform. A similar trend was also observed for intra chip reproducibility (**Figure 5B**), where the ECL signals of three dual spirals were compared across all three individual devices, showing high repeatability in device fabrication. Therefore, these results indicate a significant enhancement in ECL signal intensity through the application of a negative bias at the GE, further highlighting the spiral GC design’s capability to generate highly reproducible and strong ECL emissions from luminol oxidation products. This is in good agreement with the results of our electrochemical studies concerning electrode widths and gaps. This spiral design GC configuration, with its precise and highly reproducible manufacturing capabilities, enables the simultaneous generation of a higher density of ROSs, which act as co-reactants to amplify the ECL signal, emphasizing a key advantage of the micro spiral GC design in achieving high sensitivity on microscale sensing platforms.

Moreover, we studied the ECL response of luminol at different gap and width configurations. In this study, all devices were fabricated with a spiral structure within a 1 × 1 mm^2^ area. The results summarised in **Figure 5C** and **D**. To investigate the effect of gap size, we tested four sensor designs: W2G2, W2G5, W2G10, and W2G20. The results showed a decrease in the ECL response as the gap size increased from 2 to 20 µm (**Figure 5C**). This decline can be attributed to two factors: (i) increased gap size enhances the diffusion of products from the CE and GE, reducing their local concentration, and (ii) larger gaps result in shorter electrode lengths, thereby decreasing the electrode surface area. Among the tested designs, W2G2 exhibited the highest ECL signal, as it had both the largest surface area and the smallest gap. The effect of four different widths (W2G10, W5G10, W10G10, and W20G10) on ECL signal was also studied, as shown in **Figure 5D** and the response remained similar among different widths, well aligned with the redox cycling results (**Figure S2**). This finding emphasizes that ROSs generation and diffusion are key factors in enhancing the ECL signal of luminol with counteracting effects, greater ROSs generation enhances the signal, while increased diffusion diminishes it.

### 3.4. Possible ECL Mechanism

To investigate the possible ECL mechanism pathway for luminol-oxygen system on entwined spiral microelectrodes, we also performed two experiments to find out the role of oxygen and luminol. We degassed oxygen and saturated luminol solution using N_2_ gas. **Figure 6A** presents the ECL signal of luminol at W2G2 which decreased significantly by removing dissolved oxygen from solution, strongly suggesting the participation of dissolved oxygen for the ECL signal enhancement. The small signal observed was due to the higly efficent diffuison process of oxygen back into the solution from the atmosphere in the time between degassing and measurment. Another study was undertaken to understand the effect of luminol concentration on the ECL signal at W2G2 device. As shown in Fig.6B, the ECL signal of luminol was recorded at various concentrations of luminol ranging from 1 µM to 2 mM with and without applying the potential of -0.6 V to the GE. In the GC setup, the ECL signal of luminol was detectable at concentration of 1 µM and enhanced by increasing the concentration of luminol, levelled off at 1mM (**Figure S3-A**.). In contrast, the ECL signal without using the GE was only detectable from 100 µM and the signal was very weak (**Figure S3-B**).

**Figure 6.**
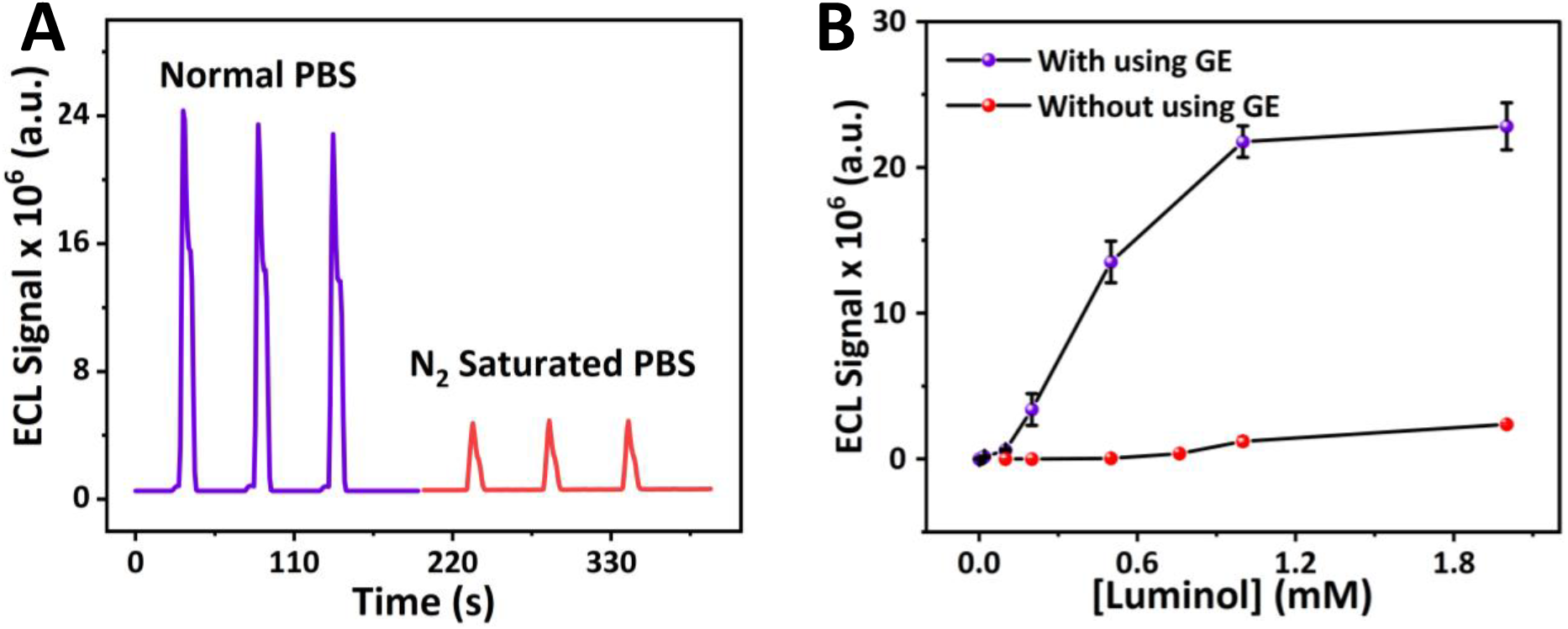
(A) the effect of Oxygen on the ECL response (B), the effect of luminol concentration on ECL response in PBS pH 7.4.

Therefore, it can be suggested as illustrated in scheme 1, with applying negative potential either with negative constant potential at the GE or by starting potential from negative potential at the CE, various reactive oxygen species (ROSs) can be produced because of the oxygen reduction reaction. At the same time by sweeping the potential in a positive direction at the CE, luminol can be oxidized to luminol radical anion and subsequent reaction between this product and ROSs can generate 3-aminophetalte in the excited state, which then emit light at 425 nm upon relaxation.^[57,58]^ Among two methods for producing ROSs, applying negative potential to a GE resulted in higher, stronger and stable ECL response which could be related to simultaneous and constant production of ROSs at the GE while potential at the CE is swept in a positive direction. However, by sweeping the potential from -0.6 to 0.8 V on the CE without using the GE, diffusion of ROSs from electrode surface toward solution and instability of the produced ROSs can decrease the number of available ROSs for subsequent CL reactions with the luminol radical anion.^[19,20,45]^ So, the two entwined micro-spiral design has tremendous potential for sensitive, coreactant free ECL analysis in a compact miniaturized platform, making ideal for portable and on-site applications.

### 3.5. Analytical applicability of coreactant free ECL platform: Trolox detection

Since ROSs are the key elements involved in the coreactant-free luminol ECL system, we further explored the potential of this ECL platform for detecting ROSs scavenger. Trolox, a water-soluble analog of vitamin E was chosen as a standard antioxidant due to its common use in evaluating antioxidant capacity in variety of samples including food, pharmaceutical and living cells. ^[59–61]^ Trolox effectively scavenges ROSs, leading to a significant quenching of the ECL signal (Scheme S1). As indicated in **Figure 7A**, increasing the concentration of Trolox in the range of 0.05 to 6 mM caused a continuous decline in the ECL intensity of luminol. As a result, at 6 mM, the ECL signal was inhibited by 99.6%. Moreover, a linear correlation was observed between the percentage of ECL signal suppression ((I_0_-I)/I_0_ ×100) and the Trolox concentration within the range of 0.05 to 2 mM and with a detection limit of 20 µM, y= 30.74 [Trolox] + 11.33, R^2^ 0.9946 (**Figure 7B**).

**Figure 7.**
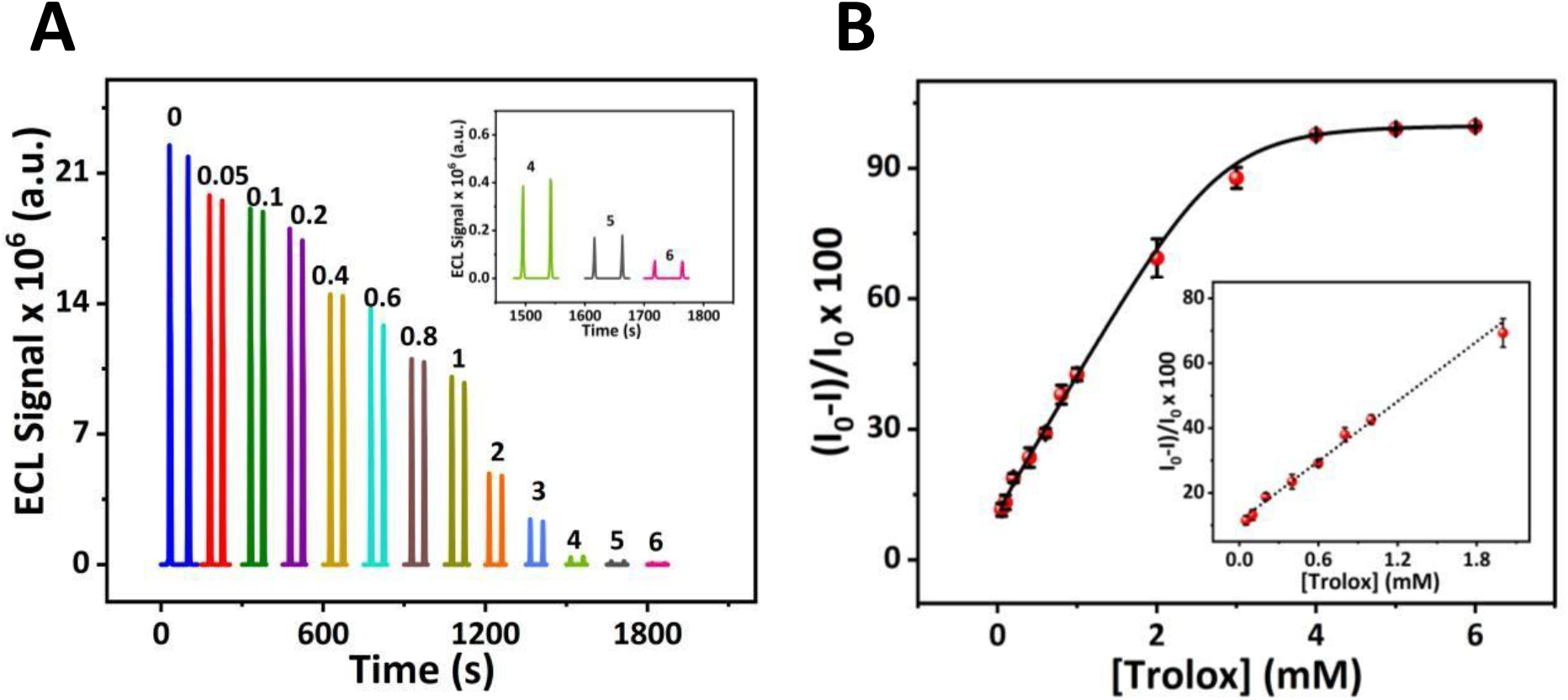
(A) ECL signal -time profile at various concentrations of Trolox and (B) calibration curve (conditions: Luminol 1mM in PBS pH 7.4, scan rate 0.1V/s).

Additionally, to evaluate the stability and restoration of the ECL signal after Trolox calibration studies, we replaced the Trolox containing luminol solution with a Trolox free luminol solution. The system displayed excellent robustness, with the ECL signal consistently returning to its initial levels after Trolox measurement, indicating potential for multiple-use analysis (**Figure S4**). The RSD% of 47 consecutive measurements over 1 hour was obtained 3.60%, implying highly stability and reproducibility of ECL signal for coreactant-free ECL system using this specific spiral CG configuration in small solution volumes (200 µL). These results indicate the reusability of this ECL platform for oxidative stress monitoring.

### 3.6. Analytical applicability of coreactant free ECL platform: hydrogen peroxide detection

We also examined the ECL response of luminol in the presence of hydrogen peroxide, both with and without applying -0.6 V to the GE. **Figure 8A** shows the ECL signal in the presence of H_2_O_2_ in the range of 100 nM to 5 mM. The ECL signal-time profiles are presented in **Figure S5** for both measurements. The ECL signal exhibited a linear relationship in the range of 10 µM to 500 µM H_2_O_2_ (y=38.66 x+1.79, R^2^ = 0.987) with a detection limit of 3.88 µM, reaching a plateau beyond this concentration when using only the CE. By applying a negative potential to the GE, the ECL signal of luminol intensifies with increasing the concentration of H_2_O_2_ from 100 nM to 500 µM and reached a plateau after that. It also showed a linear range from 10 µM to 500 µM, with a detection limit of 7.3 µM (y = 21.77 x + 3.99, R^2^ = 0.982). Furthermore, this study revealed that the enhancement effect of the entwined spiral GC setup in the luminol-O_2_ system was equivalent to that observed in the luminol-H_2_O_2_ system at 500 µM H_2_O_2_. **Figure 8B** illustrates the stability and reproducibility of the ECL signal of luminol in the presence of 1 mM hydrogen peroxide with a very low RSD of 0.144% over 39 consecutive measurements. This result confirms the reliability of the luminol-H_2_O_2_ ECL system on spiral microelectrode and its potential for H_2_O_2_ dependent ECL application as well.

**Figure 8.**
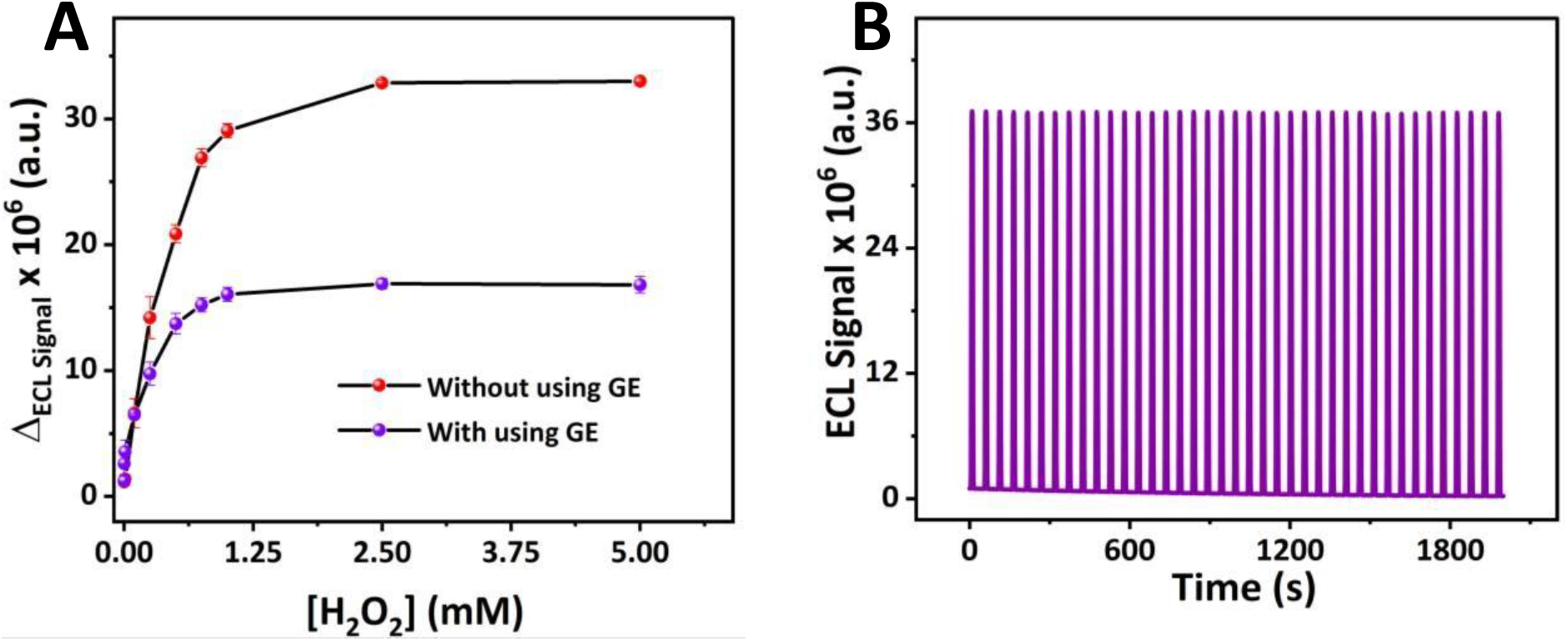
(A) the effect of hydrogen peroxide concentration on the ECL response with and without applying bias to the GE (B), stability and reproducibility of the ECL signal of luminol in the presence of 1 mM hydrogen peroxide at pH 7.4.

### 3.7. Analytical applicability of coreactant-free ECL platform for immunosensing

Next, for further validation of this platform for coreactant-free ECL immunosensing applications, both CEs and GEs were firstly modified with gold nanoparticles to increase surface area and then chitosan-gold-MWCNTs nanocomposite was applied on the CEs electrochemically to introduce amine functional groups for further selective attachment of antibody at the CEs. To develop immunosensors, W2G10 was chosen as this gap allowed uniform electrodeposition of the chitosan hydrogel onto the CE without contacting to the GE. **Figure S6** and **S7** present SEM images of the modified spiral microelectrodes with gold nanostructures and chitosan hydrogel nanocomposite at various magnifications respectively. As shown in **Figure S6**, a homogeneous deposition of gold nanostructures was successfully achieved by applying - 1mA for 5 seconds to all six spiral electrodes, indicating consistent and uniform gold deposition across the electrode surfaces. **Figure S7** also illustrates the formation of uniform layers of chitosan hydrogel incorporating MWCNTs, and gold nanostructures on the CEs. The presence of both MWCNTs and gold NPs contributed to the reproducible and uniform deposition of chitosan hydrogel nanocomposite on the spiral collector microelectrodes.

In addition, **Figure S8-A** shows CVs of the Fe(CN)_6_]^3−/4−^ redox probe at various modification steps. Upon gold NPs deposition, the current increased due to the enhanced conductivity and larger surface area provided by the gold nanostructures. Following the electrodeposition of the chitosan nanocomposite, the current decreased, attributed to the lower conductivity of the chitosan biopolymer. We also investigated the reproducibility of simultaneous gold deposition on six spiral electrodes. **Figure S8-B** illustrates clearly overlaid CVs for all six spiral microelectrodes at W2G10 device, implying high reproducibility in electrochemical modification of all six spiral microelectrodes.

To develop label free ECL immunoassay application of this multiplexed platform, anti-IgG was chosen as a model antibody representative of any IgG isotype, and protein A/G was employed to facilitate more site-directed attachment to the electrode surface. Our previous studies on the preparation of immunosensors using a silicon chip multiplexed platform demonstrated that an oriented approach could provide greater reproducibility and reliability in assay responses across multi-microelectrodes^[62]^. Therefore, after covalent binding of protein A/G to the amine groups of chitosan *via* NHS/EDC attachment chemistry, the anti-IgG antibody was immobilized onto the surface through several specific binding sites available on protein A/G for the Fc region of IgG isotype antibodies, making it a universal platform for this class of antibodies. The schematic illustration of proposed biosensors and its detection pathway is depicted in **Scheme 2**. Through this biofunctionalization, the ECL biosensor was able to detect IgG protein *via* a signal-off mechanism, where the presence of the antigen hindered further electron transfer of luminol at the CE, resulting in a decrease in ECL intensity.

**Scheme 2.**
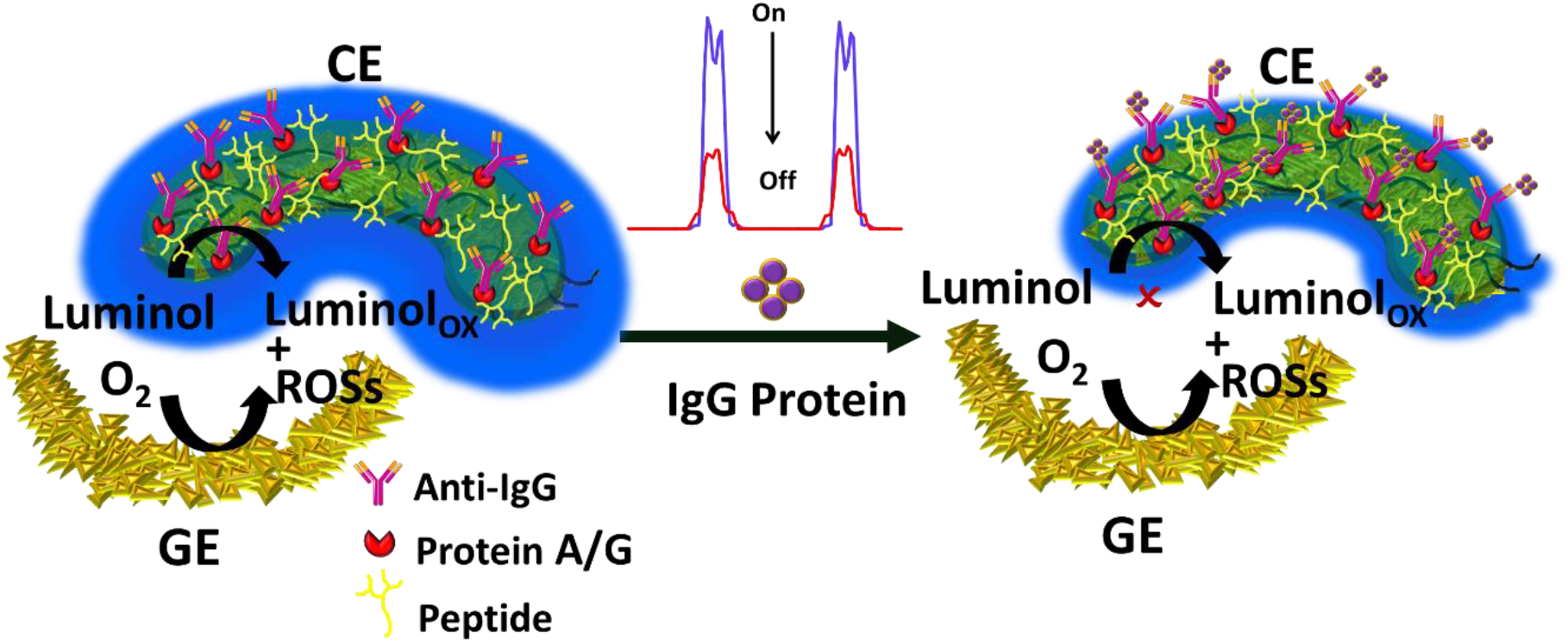
An illustration of the proposed biosensor and its On-Off ECL detection mechanism.

**Figure 9A** shows the ECL behaviour of the developed biosensor at different concentrations of IgG protein. As illustrated in the corresponding calibration curve in **Figure 9B**, the ΔI_ECL_ exhibits a linear relationship in decrease of ECL signal with the logarithmic concentration of IgG over the range of 1 fg·mL^−1^ to 100 pg·mL^−1^ (y = –0.256 log [IgG] – 0.896, R^2^ = 0.9896, n = 6). The detection limit was calculated (3σ_b_/N) as 0.8 fg·mL^−1^, clearly demonstrating the high sensitivity of this coreactant-free ECL immunosensing platform. The ECL responses of the three biosensors on a single device toward 100 fg·mL^−1^ and 100 pg·mL^−1^ IgG protein are compared in **Figure 9C**. Relative standard deviation (RSD%) values of three consecutive measurements varying from 3% to 10% were obtained among the three biosensors on each device, indicating good reproducibility and confirming the multiplexing capability of the platform. Additionally, the RSD% of biosensor responses across three independent devices varied between 2.5% and 5%, highlighting the excellent reproducibility of the biosensor fabrication process on this multiplexed platform (**Figure 9D**).

**Figure 9.**
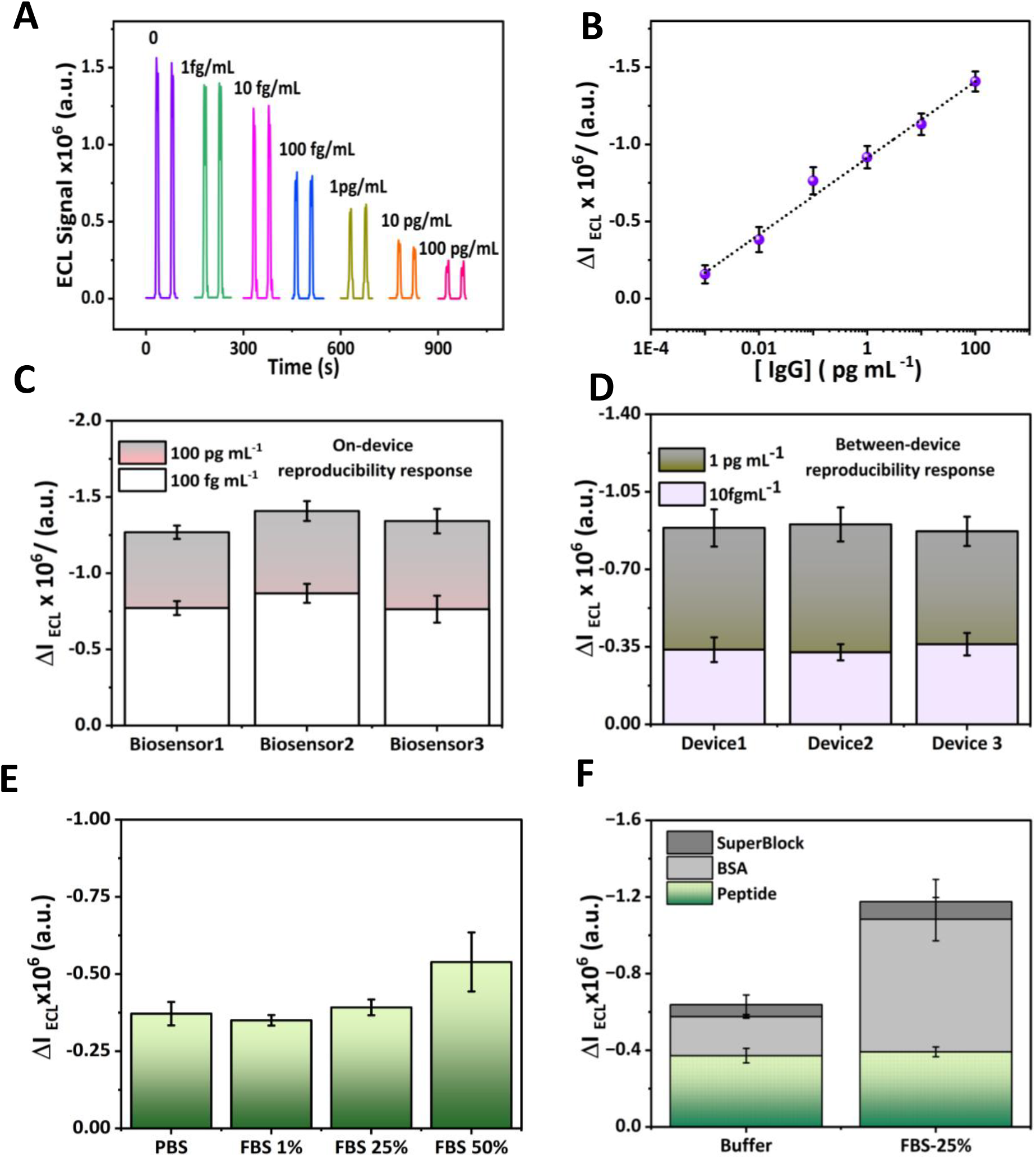
(A) ECL response of the biosensor in the presence of different concentrations of IgG protein;(B) Corresponding calibration curve for IgG detection; (C) On-device reproducibility comparison among three biosensors on one device at 100 fg mL^−1^ and 100 pg mL^−1^ IgG concentrations;(D) Between-device reproducibility comparison for 10 fg mL^−1^ and 1 pg mL^−1^ IgG; (E) Antifouling performance of the branched peptide blocking agent toward 10 fg mL^-1^ IgG in various FBS concentrations (v/v); (F) Comparison of antifouling performance among three blocking agents (branched peptide, SuperBlock, and BSA) toward 10 fg mL^−1^ IgG in 25% FBS.

To evaluate the applicability of the proposed biosensor in complex biological samples, a branched zwitterionic poly-EK peptide at a concentration of 200 µg mL^−1^ was incorporated into the biosensor design as a blocking agent. Previous study clearly demonstrated the ultralow fouling electrochemical cell detection in human serum using the branched peptide.^[63]^ **Figure 9E** depicts the antifouling performance of the branched peptide, showing the ECL response of the biosensor toward 10 fg mL^−1^ IgG at various concentrations of FBS. The results demonstrated good antifouling behaviour, with consistent responses observed even when IgG was spiked into 25% FBS samples. In normal rabbit whole serum, the IgG concentration typically ranges from 5 to 10 mg mL^−1. [64,65]^ Considering the linear range of the biosensor, the dilution factor, and the demonstrated performance in 25% FBS, the findings indicate that the biosensor can reliably detect IgG in complex biological matrices with minimal interference from fouling or nonspecific adsorption.

Moreover, we compared the antifouling performance of the branched peptide with two common blocking agents, SuperBlock and 0.1% BSA. The ECL responses of the biosensor toward 10 fg mL^−1^ IgG in 25% FBS sample were tested and compared with those obtained in PBS. As shown in **Figure 9F**, the ΔI_ECL_ in 25% FBS samples were significantly suppressed upon using SuperBlock and BSA blocking agents, with average signal reductions of 88% and 83%, respectively. Whereas ΔI_ECL_ of the branched peptide modified biosensor exhibited the same signal as in PBS under the same conditions, confirming the superior antibiofouling performance of the branched peptide. Besides, the ΔI_ECL_ for the biosensors blocked with SuperBlock and BSA were also higher compared to those using peptide blockers, suggesting that the larger size of these proteins may be less effective in reducing non-specific binding, even in PBS. In contrast, zwitterionic peptides provide steric repulsion, strong hydrophilicity, and a suitable conformation resulting in high resistance to protein adsorption in complex biological matrices as extensively reported in previous studies.^[63,66–70]^

## 4. Conclusions

In summary, we successfully designed and fabricated seven novel multiplexed sensing devices featuring entwined spiral microelectrode configurations by using highly reproducible silicon microfabrication techniques. These platforms were comprehensively evaluated for coreactant-free ECL applications, resulting in the development of a highly sensitive, miniaturized multiplexed ECL system. The unique entwined spiral design allowed to tune ECL responses and exhibited excellent sensitivity and reproducibility. The analytical performance of the platform was validated using the oxygen scavenger Trolox and hydrogen peroxide, showcasing high sensitivity, reproducibility, and reliability. Furthermore, we investigated the (bio)functionalization of the spiral microelectrodes for immunoassay applications by selectively immobilizing antibodies on the chitosan nanocomposite modified CEs. A site-selective immobilization strategy using protein A/G, combined with an antifouling branched zwitterionic peptide as a blocking agent, enabled highly sensitive and reproducible detection, both within and across biodevices, demonstrating the platform’s suitability for coreactant-free ECL immunoassays in complex biological matrices. We believe that the miniaturized design, multimodal functionality, and innovative entwined spiral electrode structure at the micrometre scale, open new avenues for remote and on-site reagent less ECL monitoring. Building on these promising results, we envision further advancing this platform by functionalizing the GEs with novel ROSs accelerator materials and the CEs with efficient luminophores to boost further the ECL signal, ultimately moving toward a fully reagent less ECL sensing system.

## Supporting Information

Supporting Information is available from the Wiley Online Library or from the author.

## ACKNOWLEDGMENT

This publication has emanated from research conducted with the financial support of the European Union Horizon Europe Programme under ‘ECL-FARM project’ (MSCA, grant agreement No. 101109381) and the European Union Horizon Europe research ‘GreenArt project’ (101060941). It has also emanated in part from research conducted with the financial support of Research Ireland and the Department of Agriculture, Food and Marine on behalf of the Government of Ireland under Grant Number [21/RC/10303_P2] – VistaMilk.

## Data Availability Statement

The data that support the findings of this study are available from the corresponding authors upon reasonable request.

