## Supporting Information for "Highly Efficient Coreactant-Free Electrochemiluminescence Sensing Platform Using Novel Microfabricated Multiplexed Entwined Spiral Microelectrodes for Point-of-Care Applications"

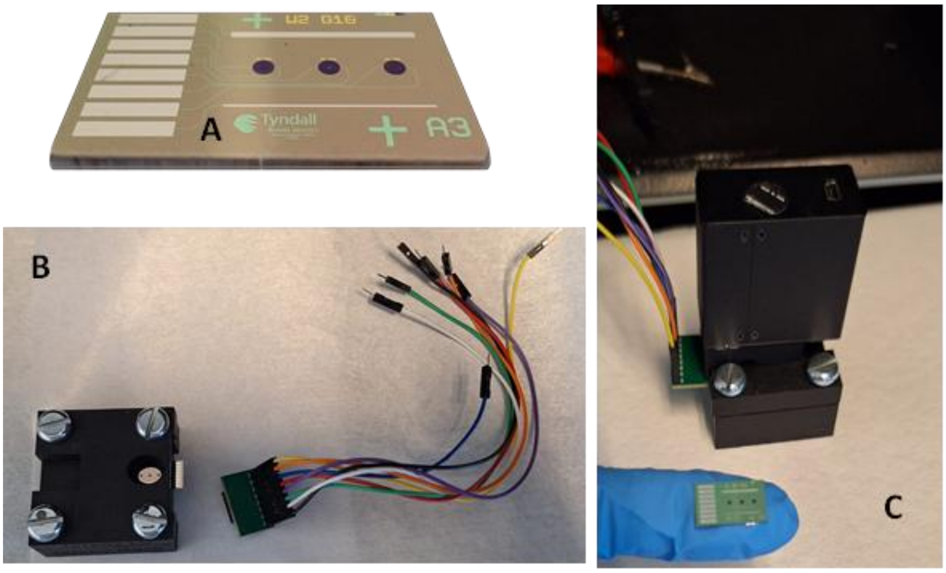


**Figure S1**. Images (A) W2G10 device, (B) cell holder with SD-connector and (D) ECL setup with PMT.

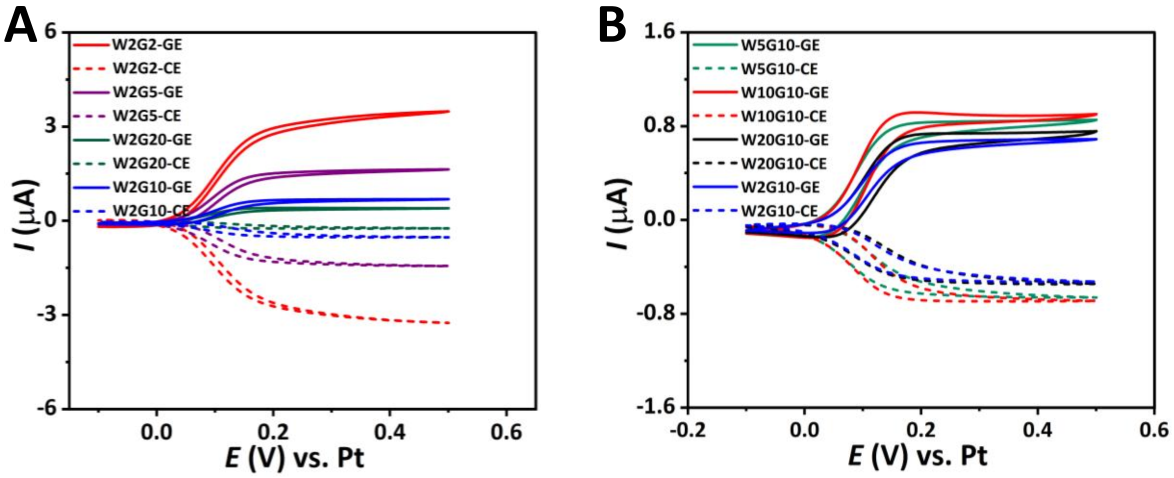


**Figure S2**. CVs of 1mM ferrocene carboxylic acid (A) at different gaps and (B) at different widths. Solids curves are CVs at GEs and dotted line are at CEs at scan rate 0.1 V/s.

**TableS1**. The effect of gaps and widths on the collection efficiency

| **Gaps** | **Collection efficiency (%)** | **Widths** | **Collection efficiency (%)** |
|---|---|---|---|
| W2G2 | 93 | W2G10 | 76 |
| W2G5 | 90 | W5G10 | 77 |
| W2G10 | 76 | W10G10 | 77 |
| W2G20 | 56 | W20G10 | 72 |

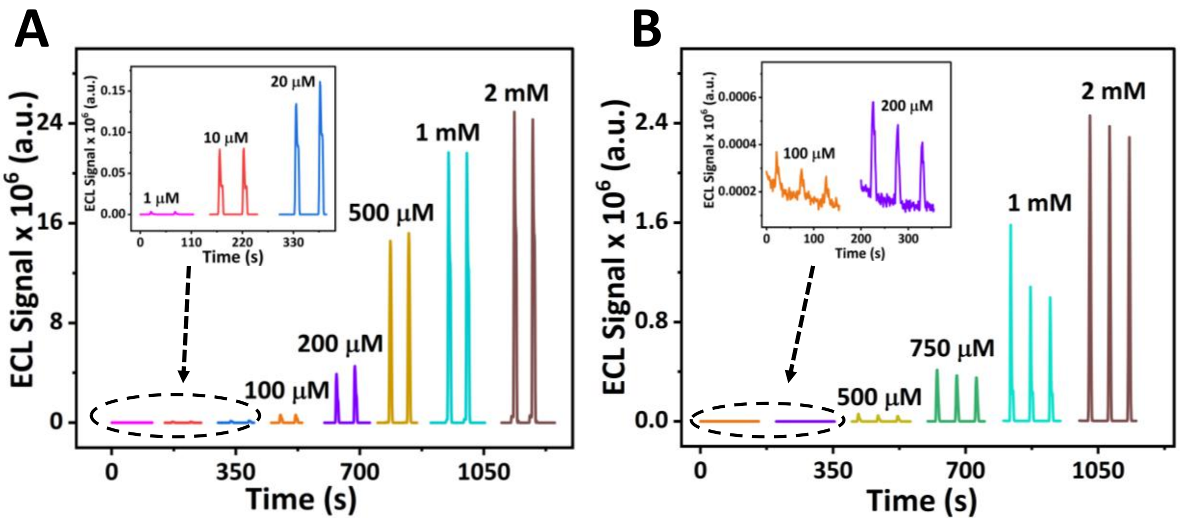


**Figure S3**. ECL signal-time profile at various concentrations of luminol (A) with and (B) without applying -0.6 V to the GE.

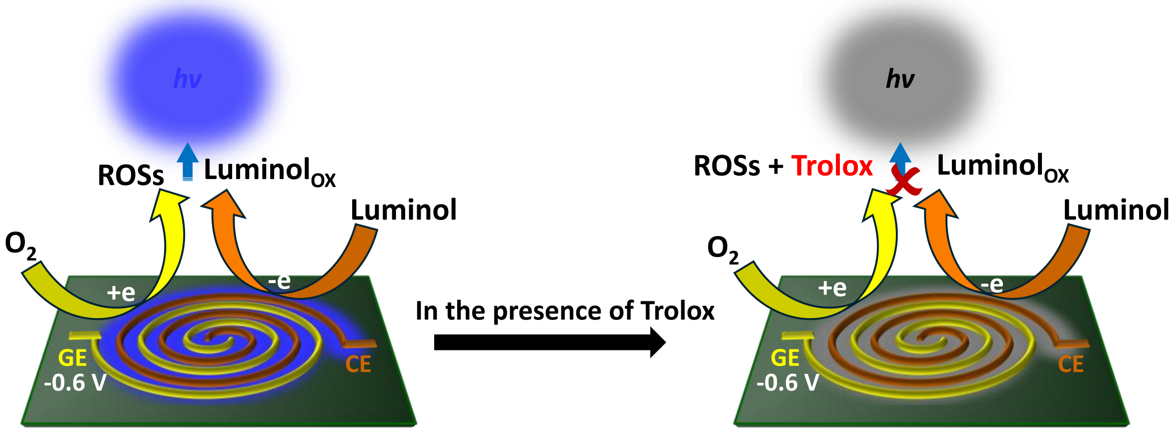


**Scheme S1.** ECL mechanism pathway of luminol at entwined spiral microelectrodes in the presence of Trolox antioxidant.

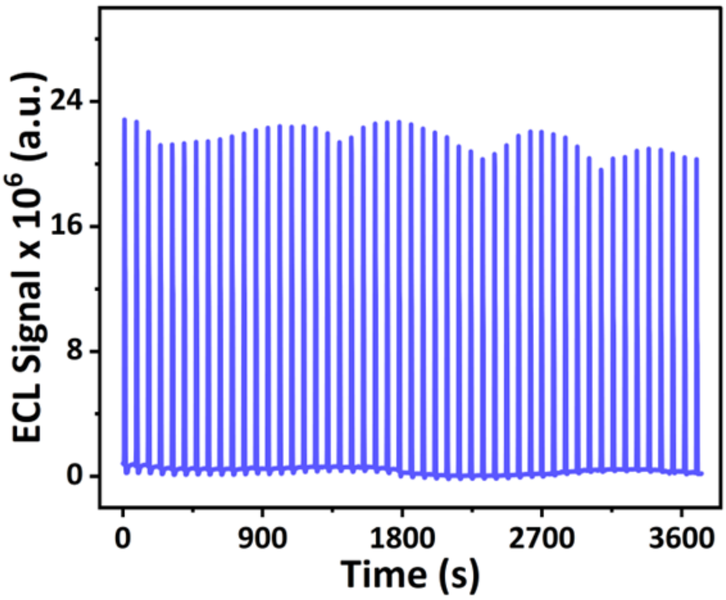

**Figure S4**. Stability test of the restored ECL Signal after Trolox measurement in PBS buffer pH 7.4 containing 1mM Luminol.

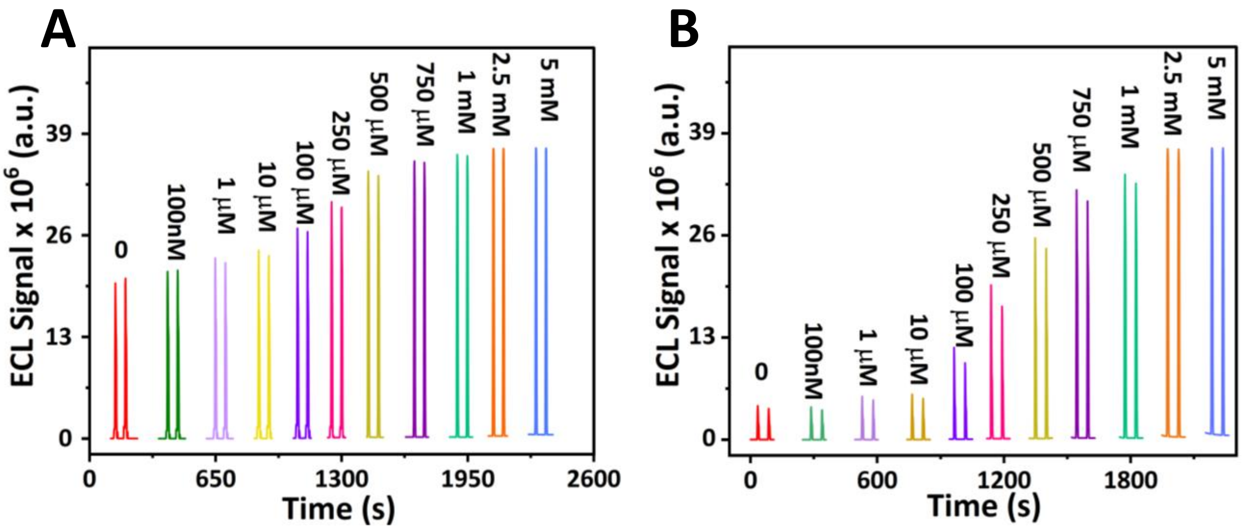


**Figure S5**. The ECL signal-time profiles of luminol 1mM in PBS pH 7.4 in the presence of different concentrations of $H_2O_2$ (A) with and (B) without applying bias to the GE.

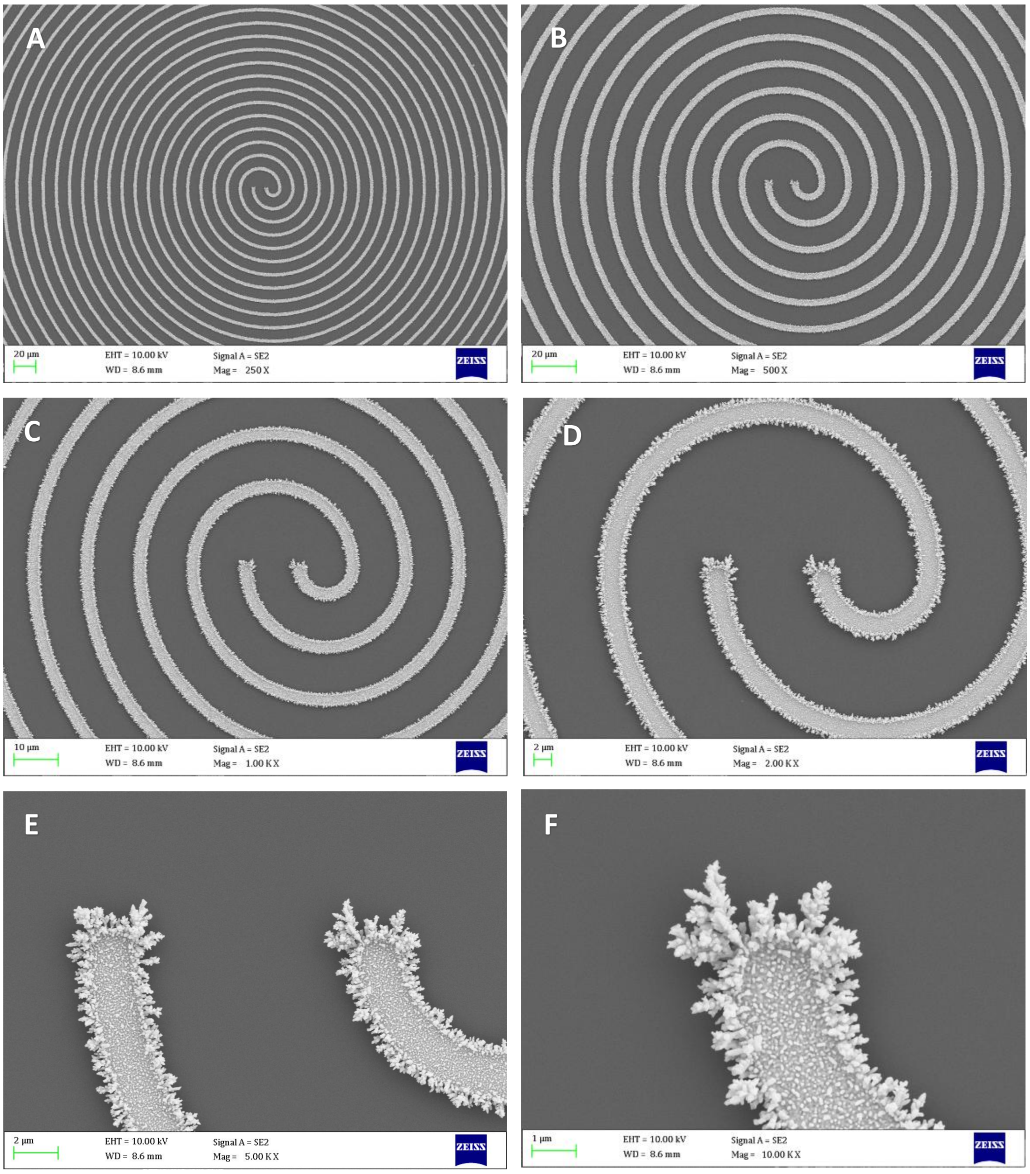


**Figure S6**. SEM images of gold nanostructured modified both CE and GE piral microelectrodes at W2G10 device at different magnifications.

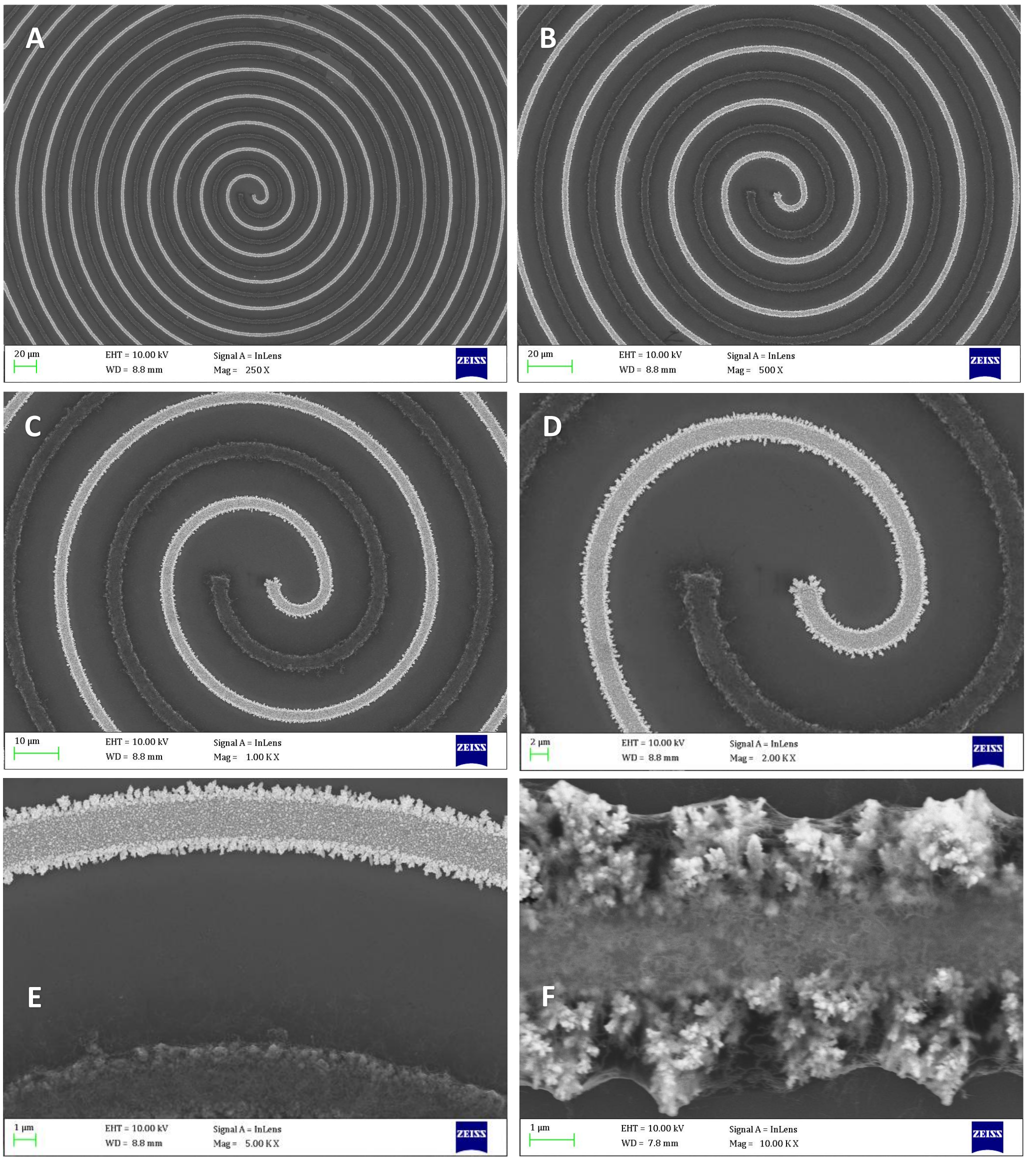


**Figure S7**. SEM images of chitosan hydrogel-MWCNTs-gold nanocomposite modified at the CE and gold nanostructured the GE on W2G10 device at different magnifications.

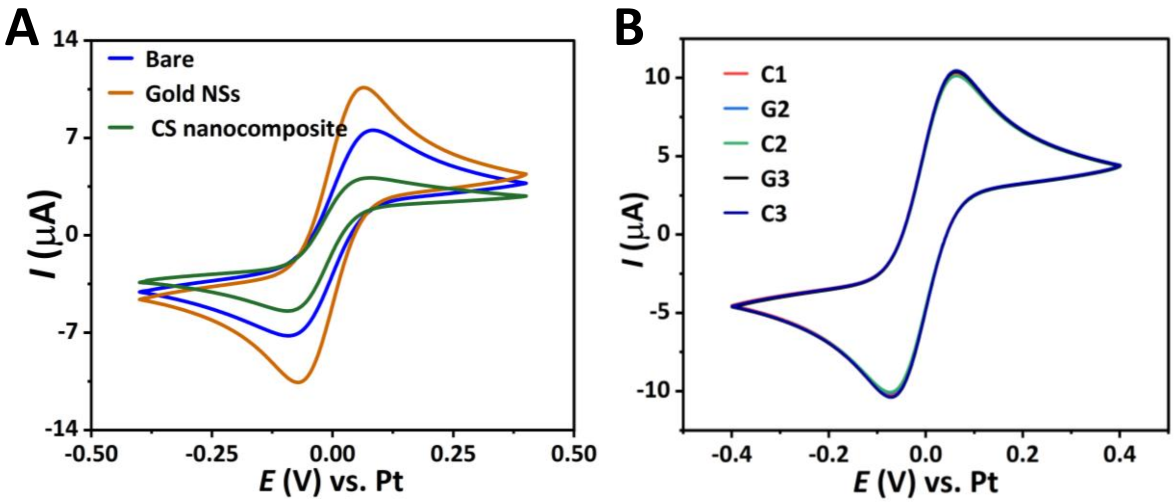


**Figure S8**. CVs of 5mM $Fe(CN)_6]^{3-/4-}$ redox probe (A) recorded at bare gold electrodes, gold NSs and chitosan nanocomposite modified electrodes, (B) at all GEs and CEs at scan rate 100 mV $s^{-1}$.
